## Supplementary material for "Identification of potential human pancreatic *α*-amylase inhibitors from natural products by molecular docking, MM/GBSA calculations, MD simulations, and ADMET analysis": Supplementary_Material.docx

Supplementary Contents

### 1. Supplementary tables:

Table S1. Ligand library of natural products

| Source of  Inhibitor | Inhibitor | Structure | IC_50_ /% of  Inhibition | Ref |
| --- | --- | --- | --- | --- |
| *Ocimum tenuiflorum* | dehydrodieugenol B | 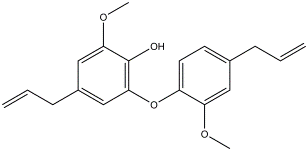 | 29.6 μM | [1] |
| Olive mill  wastes (OMW) | 1-acetoxypinoresinol | 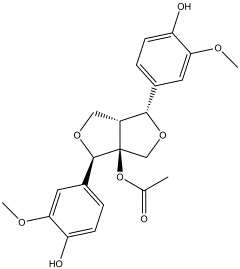 | 13.9 μM | [2] |
| *Sargassum patens* | DDBT | 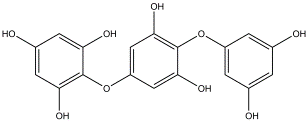 | 3.2μg/mL | [3] |
| *Humulus lupulus* | 3ʹ-geranyl  chalconaringenin | 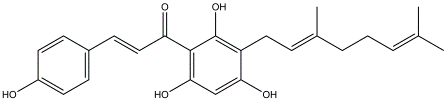 | 20.46 μM | [4] |
| *Azadirachta indica* | gedunin | 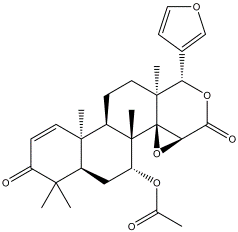 | 68.38 μM | [5] |
| *Dioscorea bulbifera* | diosgenin | 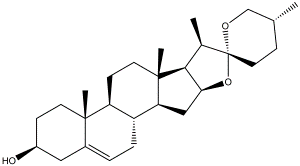 | 70.94 % | [6] |
| *Swertia chirata* | mangiferin | 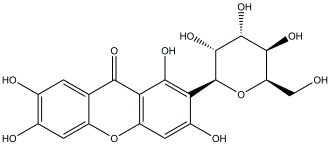 | 516.66 μM | [7] |
| *Syzygium cumini* | ursolic acid | 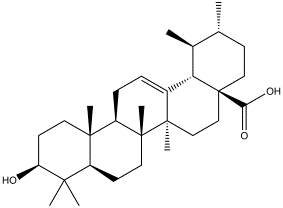 | 6.7 μg/mL | [8] |
| *Syzygium cumini* | oleanolic acid | 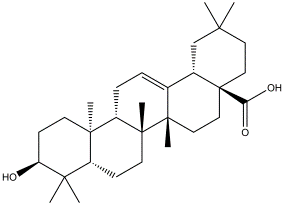 | 57.4  μg/mL | [8] |
| *Abrus precatorius* | lupenone | 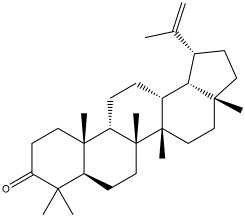 | 31 μM | [9] |
| *Psidium guajava Linn* | myricetin | 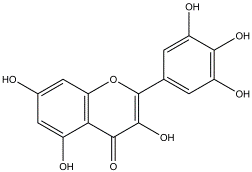 | 4.3 mM | [10] |
| *Punica granatum* | valoneic acid dilactone | 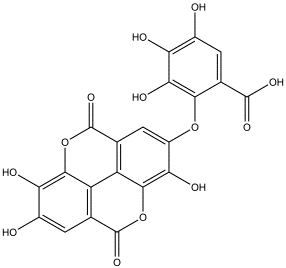 | 0.284  μg/mL | [11] |
|  | acarbose | 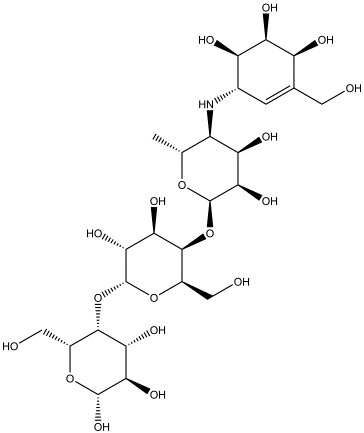 | Standard |  |
| *Passiflora ligularis* Juss | quercetin-3-O-*β*-glucoside | 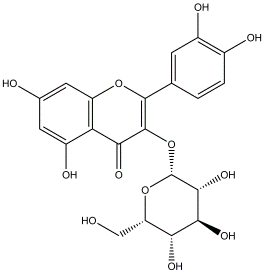 | 31.0μM | [12] |
| *Passiflora ligularis* Juss | kaempferol-3-O-*β*-glucoside | 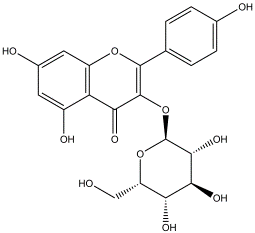 | 33.4μM | [12] |
| *Passiflora ligularis* Juss | ligularoside A | 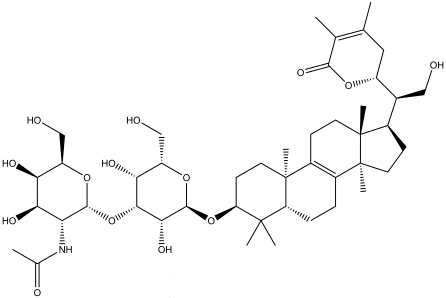 | 409.8 μM | [12] |
| *Rheum turkestanicum* | daucosterol | 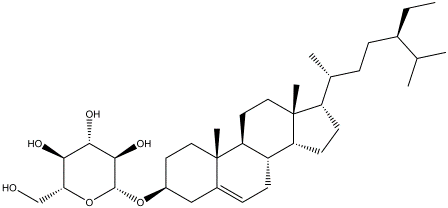 | 46.4 μM | [13] |
| *Rheum turkestanicum* | rhododendrin | 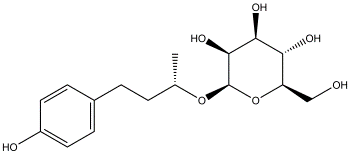 | 107.5μM | [13] |
| *Rheum turkestanicum* | emodin | 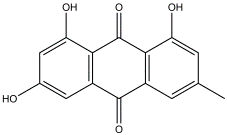 | 184.7 μM | [13] |
| *Zanthoxylum chalybeum* | chaylbemide A | 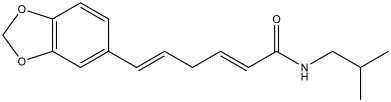 | 45.76 μM | [14] |
| *Zanthoxylum chalybeum* | chaylbemide B | 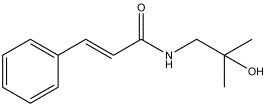 | 43.22 μM | [14] |
| *Zanthoxylum chalybeum* | chaylbemide C | 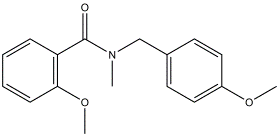 | 46.76 μM | [14] |
| *Zanthoxylum chalybeum* | trans-fagaramide | 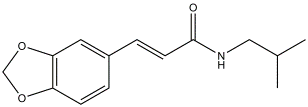 | 47.36 μM | [14] |
| *Zanthoxylum chalybeum* | skimmianine | 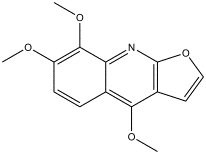 | 47.72 μM | [14] |
| *Zanthoxylum chalybeum* | norchelerythrine | 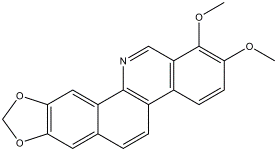 | 46.49 μM | [14] |
| *Zanthoxylum chalybeum* | sesamine | 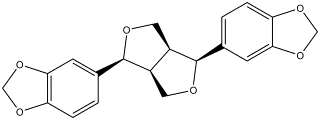 | 54.67 μM | [14] |
| *Musa acuminate* | cycloeucalenone | 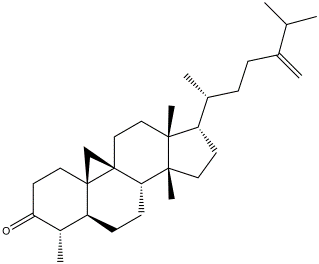 | 20.33 μM | [15] |
| *Dipterocarpus littoralis* | *α*-viniferin | 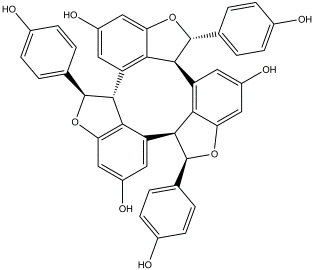 | 212.79 μg/mL | [16] |
| *Melicope latifolia* | halfordin | 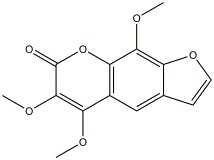 | 197.53 μM | [17] |
| *Melicope latifolia* | *β*-sitosterol | 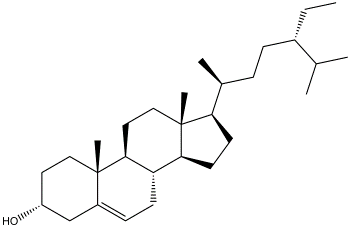 | 372.31 μM | [17] |
| *Newbouldia laevis* | newboulaside A | 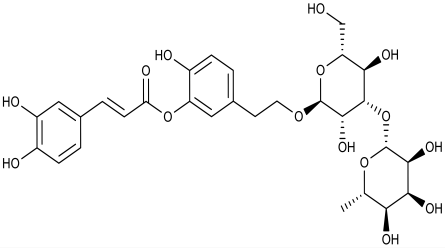 | 4.95 µg/mL | [18] |
| *Newbouldia laevis* | newboulaside B | 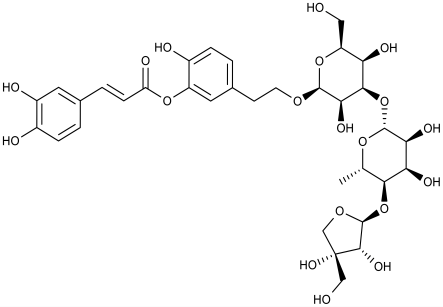 | 4.44 µg/mL | [18] |
| *Camellia sasanqua* | sasastilboside A | 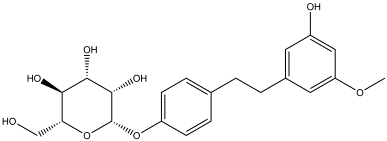 | 53.7 μM | [19] |
| *Salvia virgata* | chrysoeriol | 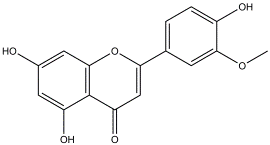 | 1.27 mM | [20] |
| *Oryza sativa* cv | momilactone A | 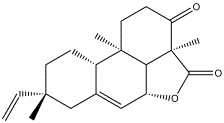 | 266.68 μg/mL | [21] |
| *Oryza sativa* cv | momilactone B | 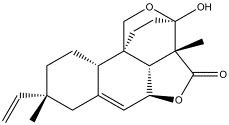 | 146.85  μg/mL | [21] |
| *Setosphaeria rostrata* | rostratazine B | 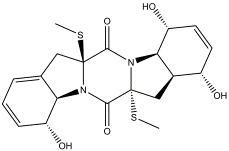 | 578 μM | [22] |
| *Physalis peruviana* | peruviose D | 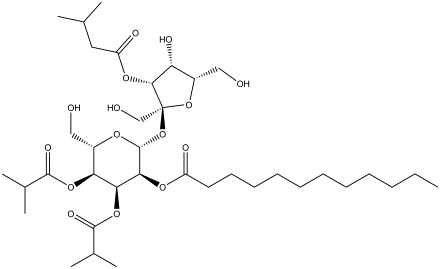 | 84.8% | [23] |
| *Physalis peruviana* | peruviose B | 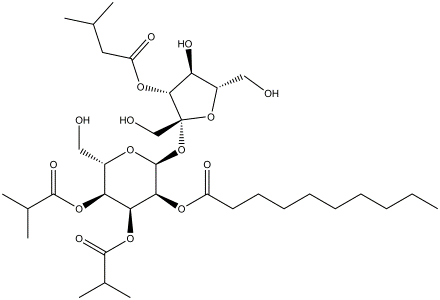 | 78.2% | [23] |
| *Physalis peruviana* | peruviose E | 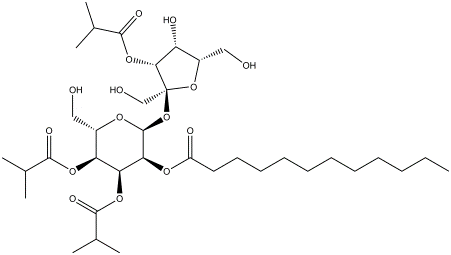 | 61% | [23] |

Key

DDBT	2-(4-(3,5-dihydroxyphenoxy)-3,5-dihydroxyphenoxy) benzene-1,3,5-triol

Table S2. Human pancreatic *α*-amylase

| Name of Target Structure | PDB ID | Resolution (Å) | Reference |
| --- | --- | --- | --- |
| Human Pancreatic *α*-Amylase | 2QV4 | 1.97 | [24] |

Table S3. SiteMap result

| Title | SiteScore | Dscore | Volume | Balance | Residues |
| --- | --- | --- | --- | --- | --- |
| Sitemap_site_1 | 0.985 | 1.01 | 341.285 | 0.751 | 58,59,62,63,101,151,  162,165,195,197,198,  200,201,233,234,235,  299,300,305 |

Table S4. ADMET properties of the acarbose and newboulaside B by QikProp

| Compound | Mol. Wt. | DonorHB | AcceptHB | QPlogPo/w | QPlogS | QPlogKhsa |
| --- | --- | --- | --- | --- | --- | --- |
| Recommended Range | 130.0  –  725.0 | 0.0  –  6.0 | 2.0  –  20.0 | −2.0  –  6.5 | −6.5  –  0.5 | −1.5  –  1.5 |
| acarbose | 645.6 | 14 | 32.1 | -7.254 | 0.739 | -2.601 |
| newboulaside B | 756.7 | 11 | 27.6 | -2.995 | -2.52 | -2.173 |

Key[25]

Donor HB	- Number of hydrogen bonds that would be donated

Accept HB	- Number of hydrogen bonds that would be accepted

QPlogPo/w	- Octanol/water partition coefficient

QPlogS	- Aqueous solubility

QPlogkhsa	- binding to human serum albumin

Table S5. ADMET properties of newboulaside B and acarbose by admetSAR

| Property | newboulaside B | | acarbose | |
| --- | --- | --- | --- | --- |
| ADMET Profile | Value | Probability | Value | Probability |
| Human Intestinal Absorption | + | 0.6701 | - | 0.9623 |
| OATP2B1 inhibitor | - | 1 | - | 0.8642 |
| OATP1B1 inhibitor | + | 0.8726 | + | 0.85 |
| OATP1B3 inhibitor | + | 0.9568 | + | 0.9497 |
| MATE1 inhibitor | - | 0.84 | - | 1 |
| OCT2 inhibitor | - | 0.875 | - | 0.95 |
| BSEP inhibitor | + | 0.7961 | - | 0.8825 |
| CYP3A4 inhibition | - | 0.8812 | - | 0.9919 |
| CYP2C9 inhibition | - | 0.8041 | - | 0.8639 |
| CYP2C19 inhibition | - | 0.8473 | - | 0.8109 |
| CYP2D6 inhibition | - | 0.8883 | - | 0.8944 |
| CYP1A2 inhibition | - | 0.8622 | - | 0.852 |
| Carcinogenicity (binary) | - | 0.9571 | - | 0.9857 |
| Ames mutagenesis | - | 0.61 | - | 0.53 |
| Skin sensitization | - | 0.8197 | - | 0.8681 |
| Mitochondrial toxicity | - | 0.525 | + | 0.775 |
| Nephrotoxicity | - | 0.8165 | - | 0.8763 |
| **ADMET profile** | **Value** | **Unit** | **Value** | **Unit** |
| Acute Oral Toxicity | 1.467 | log(1/  (mol/kg)) | 0.944 | log (1/  (mol/kg)) |

Table S6. Toxicity profile of newboulaside B and acarbose by ProTox-II

| Property | newboulaside B | | acarbose | |
| --- | --- | --- | --- | --- |
| Toxicity Class | V | | VI | |
| **Target** | **Prediction** | **Probability** | **Prediction** | **Probability** |
| Hepatotoxicity | Inactive | 0.84 | Active | 0.57 |
| Carcinogenicity | Inactive | 0.81 | Inactive | 0.82 |
| Mutagenicity | Inactive | 0.78 | Inactive | 0.72 |
| Cytotoxicity | Inactive | 0.77 | Inactive | 0.68 |

Table S7. ADMET properties of newboulaside B (1) and acarbose (2) by pkCSM

| Property | Model Name | **1** | **2** |  | Unit |
| --- | --- | --- | --- | --- | --- |
| Absorption | P-glycoprotein I inhibitor | No | No |  | Categorical (Yes/No) |
| Absorption | P-glycoprotein II inhibitor | No | No |  | Categorical (Yes/No) |
| Distribution | BBB permeability | -2.762 | -2.449 |  | Numeric (log BB) |
| Distribution | CNS permeability | -5.747 | -7.177 |  | Numeric (log PS) |
| Toxicity | hERG I inhibitor | No | No |  | Categorical (Yes/No) |
| Toxicity | hERG II inhibitor | No | Yes |  | Categorical (Yes/No) |

Table S8. ADMET properties of newboulaside B and acarbose by SwissADME

| **Property** | | | **newboulaside B** | **acarbose** | | |
| --- | --- | --- | --- | --- | --- | --- |
| **Pharmacokinetics** | | | | | | |
| GI absorption | | Low | | | Low | |
| BBB permeant | | No | | | No | |
| P-gp substrate | | No | | | Yes | |
| Log Kp (skin permeation) | | -12.08 cm/s | | | -16.30 cm/s | |
| **Drug Likeness** | | | | | | |
| Lipinski | No; 3 violations: MW>500, NorO>10, NHorOH>5 | | | | | No; 3 violations: MW>500, NorO>10, NHorOH>5 |

### 2. Supplementary figures:

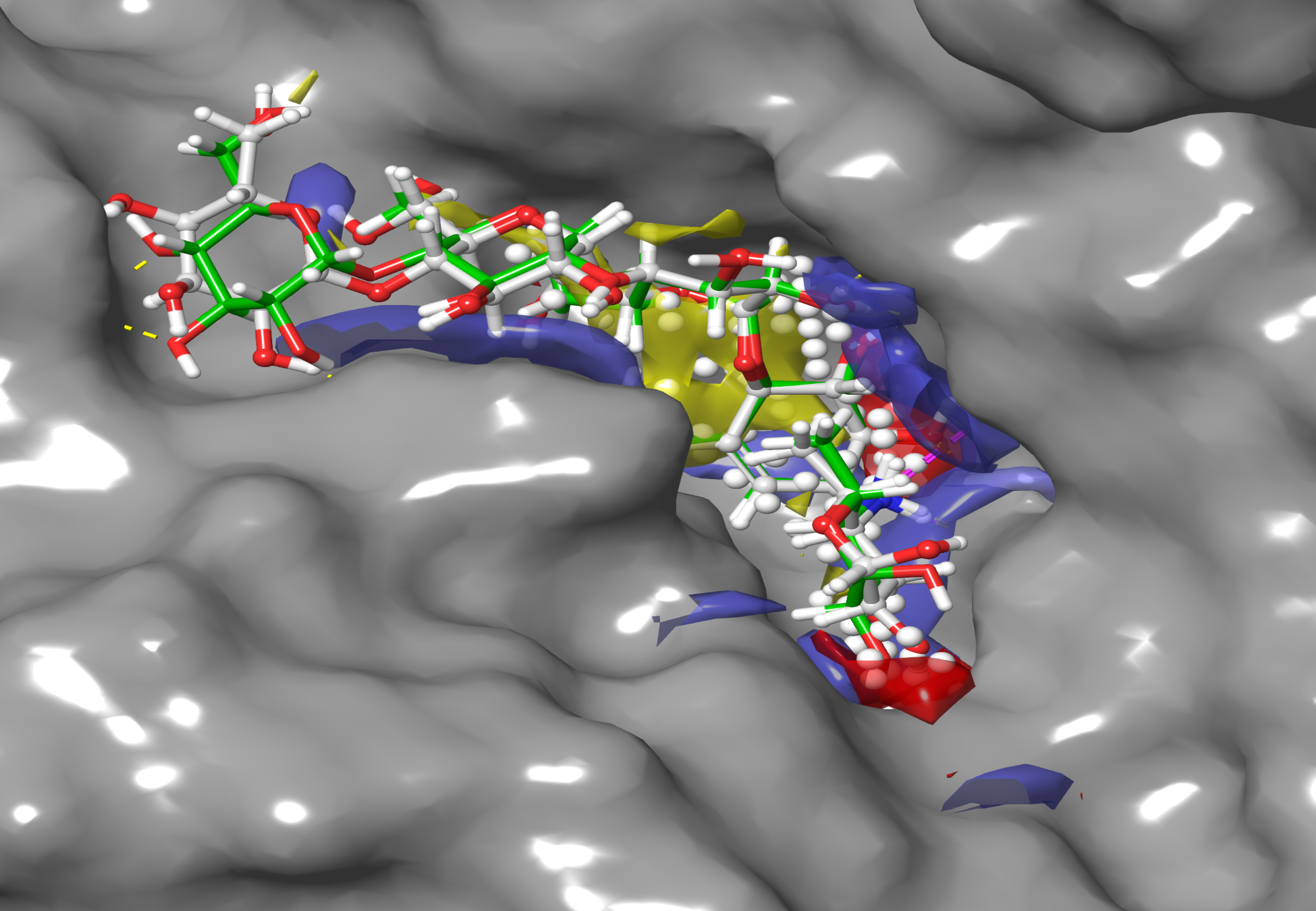

Fig S1. Pictorial representation of HPA with cognate ligand (green) and re-docked cognate ligand (white). The yellow and blue patches represent the SiteMap application view.

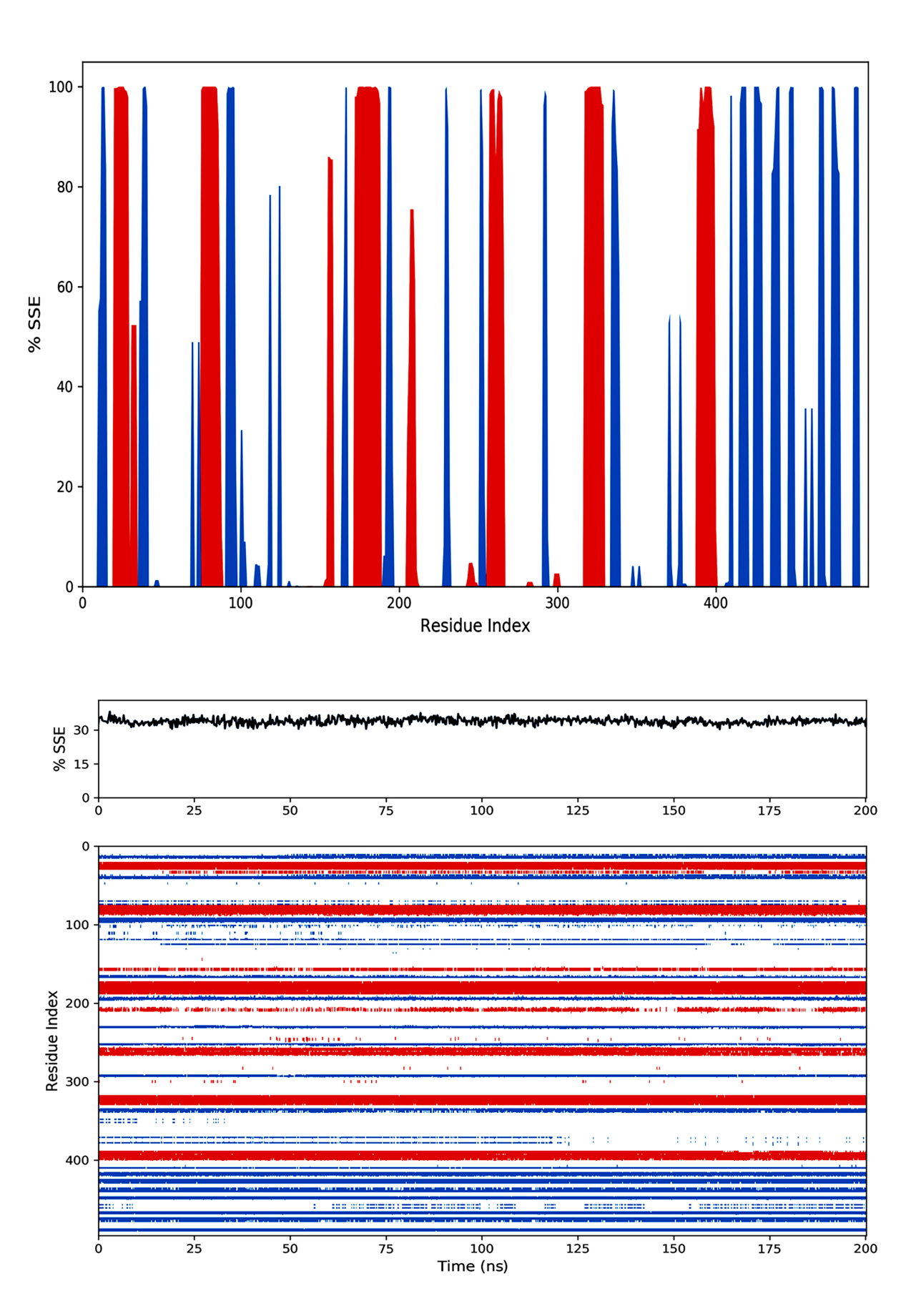

Fig S2. Analysis of secondary structure element (SSE) of HPA after MD simulation: The strand, helix, and loop region of HPA are represented as blue, red, and white, respectively.

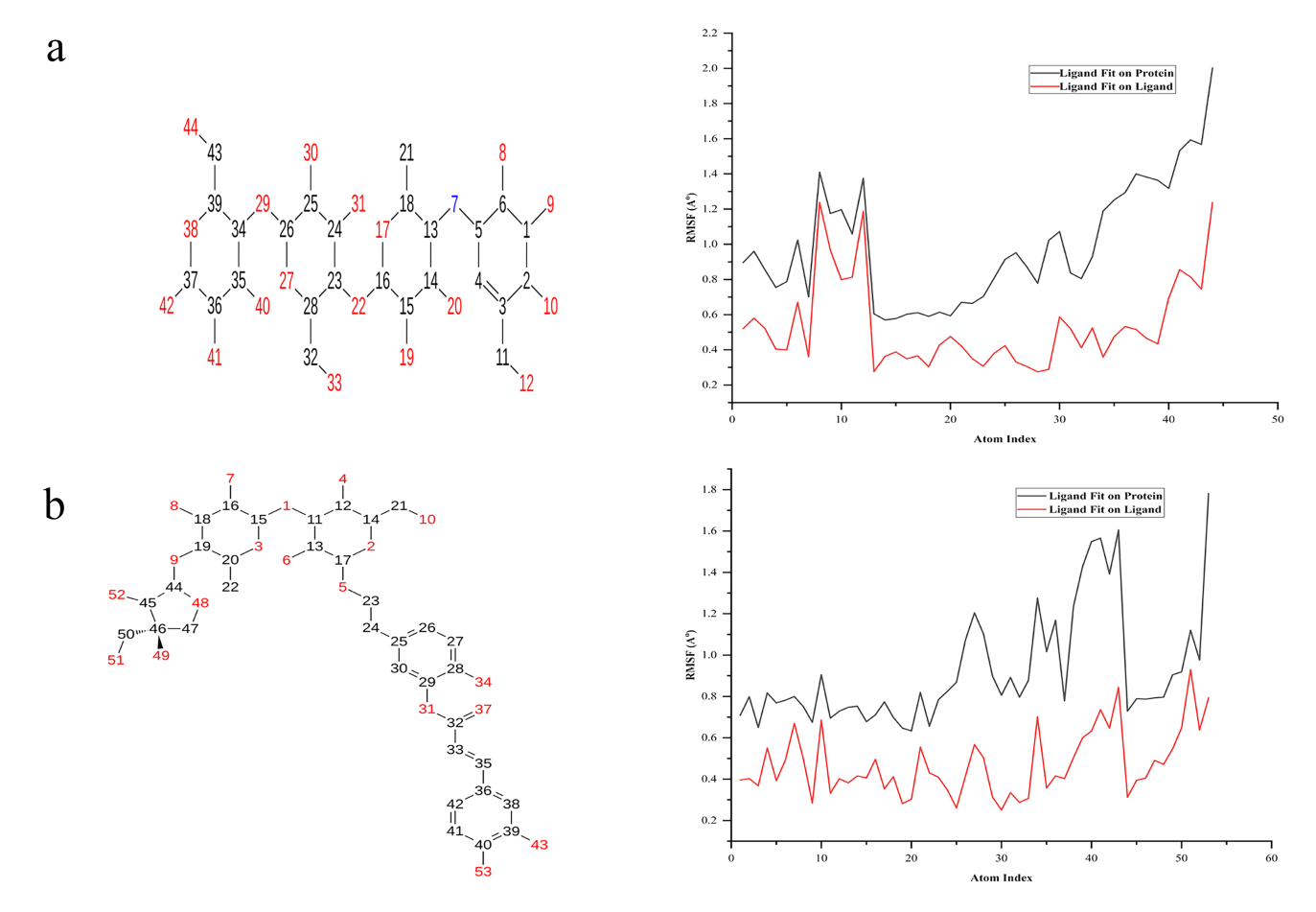

Fig S3. RMSF of inhibitor’s atoms after MD simulation: (a) acarbose and (b) newboulaside B

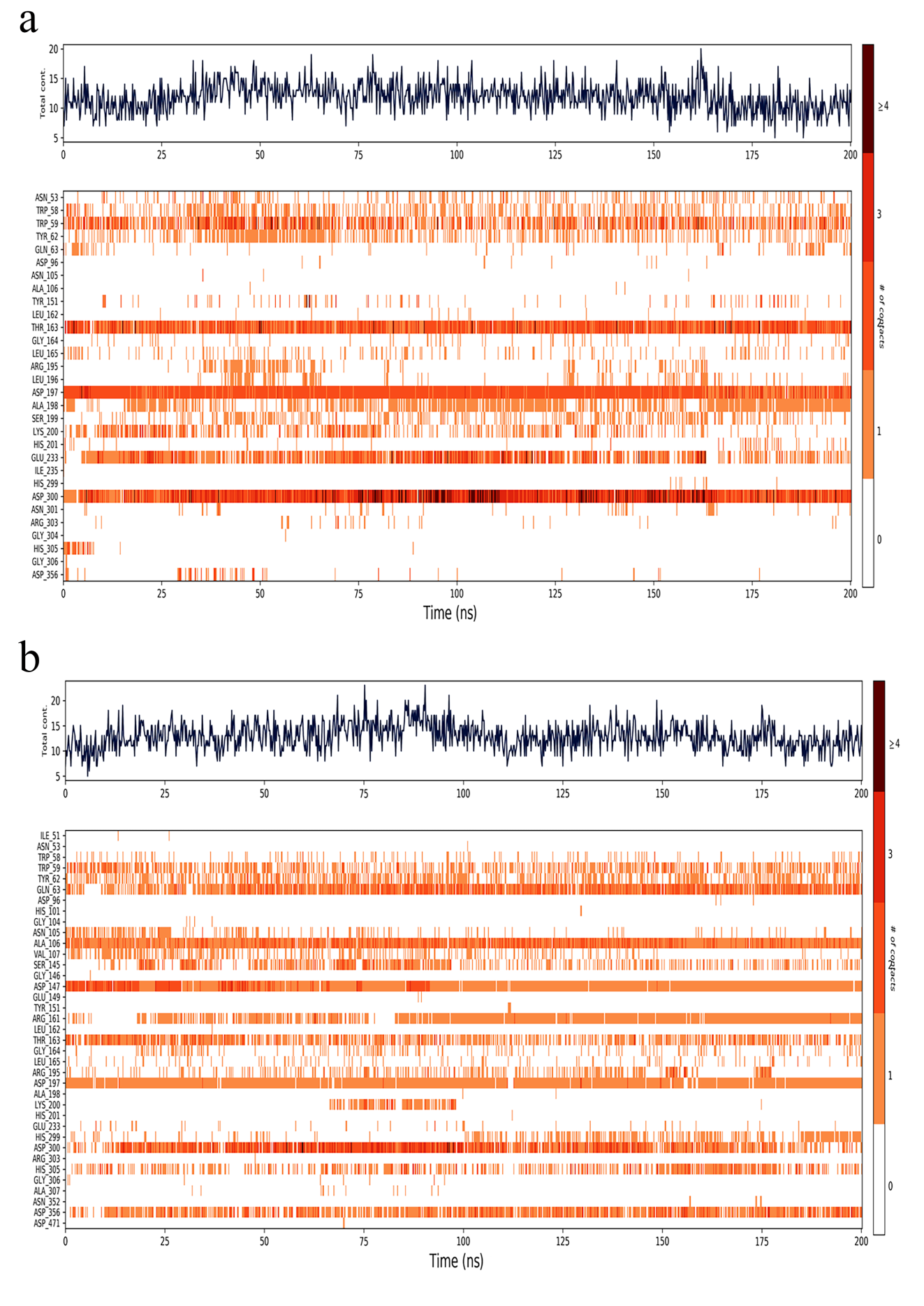

Fig S4. Timeline representation of protein-ligand contact of MD simulation trajectory: (a) HPA acarbose complex (b) HPA newboulaside B complex

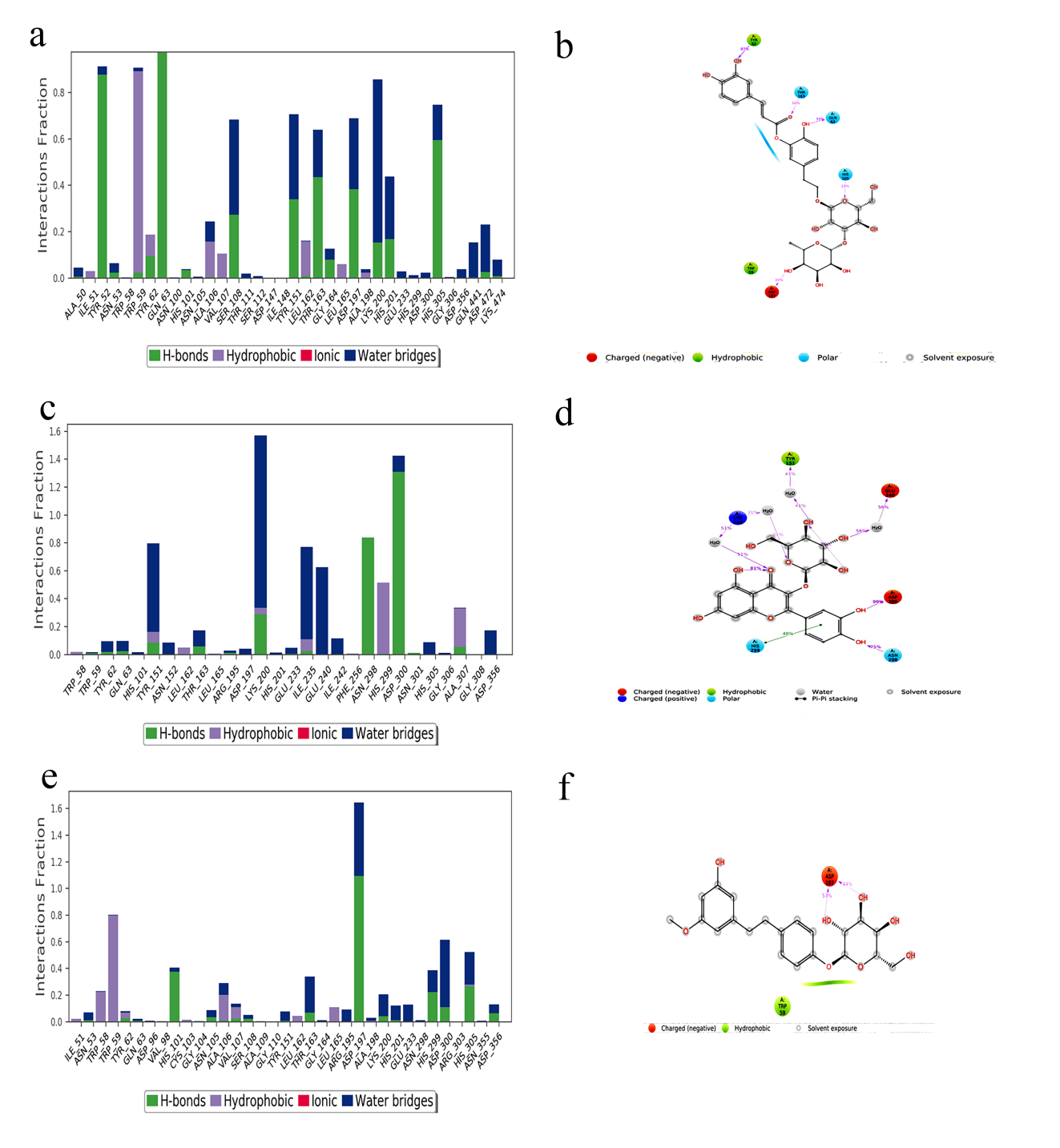

Fig S5. Analysis of inhibitor’s molecular interaction and type of contact with HPA after MD simulations: Normalized stacked bar chart of HPA binding site residues interacting with (a) newboulaside A, (c) quercetin-3-O-*β*-glucoside, and (e) sasastilboside A via hydrogen bonds, hydrophobic and ionic interactions and water bridges. Detailed schematic interaction of (b) newboulaside A, (d) quercetin-3-O-*β*-glucoside, and (f) sasastilboside A atoms with the binding site residues of HPA. Interactions happening more than 30% of the simulation times are shown.

https://doi.org/10.1023/B:JCAM.0000021861.31978.da
